# Demonstration of resistance to satyrization behavior in *Aedes aegypti* (Linnaeus) from La Réunion island

**DOI:** 10.1101/2020.02.10.942839

**Authors:** Hamidou Maïga, Jérémie R. L. Gilles, Rosemary Susan Lees, Hanano Yamada, Jérémy Bouyer

## Abstract

*Aedes aegypti* and *Aedes albopictus* are competent vectors of arboviruses such as dengue and chikungunya viruses which co-exist in some areas including La Réunion island. A kind of reproductive interference called satyrization has been described in sympatric species where a dominant species tends to control the spread of the other species. Here, we investigated satyrization in laboratory experiments to provide insights on the potential impact on *Ae. aegypti* of a control campaign including a sterile insect technique component against *Ae. albopictus*. Different mating crosses were used to test sympatric, conspecific-interspecific and allopatric effects of irradiated and non-irradiated male *Ae. albopictus* on female *Ae. aegypti*, including in a situation of skewed male ratio. Our results suggest that there was only a low level of satyrization between sympatric populations of *Ae. aegypti* and *Ae. albopictus* colonised from La Réunion island. A male *Ae. albopictus* to female *Ae. aegypti* ratio of 3:1 did not increase the level of satyrization. Female *Ae. aegypti* previously mated to male *Ae. albopictus* were not prevented from being inseminated by conspecific male *Ae. aegypti*. A satyrization effect was not seen between allopatric *Ae. albopictus* and *Ae. aegypti* strains from La Réunion Island either. The tested *Ae. aegypti* strain from La Réunion island has therefore developed full resistance to satyrization and so releasing sterile male *Ae. albopictus* may not suppress *Ae. aegypti* populations if an overflooding of irradiated male *Ae. albopictus* leads to similar results. The management strategy of two competent species in a sympatric area is discussed.

## INTRODUCTION

*Aedes albopictus* (Skuse), the Asian tiger mosquito, has been invasive in many parts of the world since the 80s (Benedict et al. 2007). *Aedes aegypti* (Linnaeus), also called the yellow fever mosquito and originating from Africa, is a highly invasive, medically important mosquito species which has received a considerable increase in attention after being linked to the Zika outbreak in Brazil in 2015 (de Araújo et al. 2016). Both species transmit several arboviral diseases including dengue, chikungunya, yellow fever and Zika (Levy Blitchtein et al. 2016). Dengue alone is estimated to infect 390 million people per year, causing 96 million cases with clinical manifestations (Bhatt et al. 2013). *Aedes albopictus* (Skuse) and *Ae. aegypti* are the most important vectors of the epidemic forms of dengue and chikungunya viruses to humans (Kyle & Harris, 2008; Paupy et al. 2010). *Aedes albopictus* is also responsible for the major chikungunya outbreak in the islands of the South-West Indian Ocean, including La Réunion island (an overseas department of France) between 2005 and 2007 (Delatte et al. 2008) and for the more recent dengue outbreaks according to the local health authority (Agence Régionale de Santé Océan Indien) and the regional office of Santé publique France on La Réunion island (WHO 2019).

The heavy reliance on insecticides to control adult *Aedes* mosquitoes, especially during disease outbreaks, has led to the emergence of widespread resistance to these chemicals, making traditional control strategies insufficient to achieve adequate reduction of vector populations. The use of insecticides is also inefficient against these container-breeding mosquito species with disseminated larval habitats. Therefore, complementary vector control methods are needed to enhance existing efforts (WHO 2017). Amongst those being advocated is the sterile insect technique (SIT), a species-specific and environmentally-friendly pest population control method which relies on maintaining a continuous production and repeated release of over-flooding numbers of sterile males (Knipling, 1959) that can outcompete their wild counterparts within the target area (Dyck et al. 2005) and induce sterility in wild females. A feasibility assessment of an area-wide integrated pest management (AW-IPM) program with an SIT component is ongoing on La Réunion island, where *Ae. albopictus* co-occurs with *Ae. aegypti*. The first releases of sterile male *Ae. albopictus* mosquitoes to study their behavior in urban areas were recently authorized by a prefectural order to the Institute for Research for Development (IRD) (Arrêté No 2019-2213/SG/DRECV). To successfully and cost-effectively apply the SIT in an area, it is recommended to target one species at a time (Dyck et al. 2005; Alphey et al. 2010). For example, where *Ae. aegypti* and *Ae. albopictus* are sympatric and are both competent vectors of human viruses, one would consider the best approach to guarantee successful suppression or elimination of both species. The best scenario would be the release of one species which was able to also readily mate with and induce sterility in the other species (Honma et al. 2019). This phenomenon is known as reproductive interference or satyrization, in which males of one species mate with and sterilize females of another species, and in this way contribute to its displacement from the shared area (Ribeiro 1988). Reproductive interference has been described in sympatric species including ticks in Mozambique (Bournez et al. 2015).

Satyrization was proposed as a possible mechanism for the displacement of *Ae. aegypti* by *Ae. albopictus* in Florida in the late 80s (Nasci et al. 1989; Tripet et al. 2011). Satyrisation and other factors such as larval competition, climate and socioeconomic factors have influenced the distribution dynamics of the two species worldwide. A rapid decline in *Ae. aegypti* in the south eastern USA and Bermuda, for example, was associated with the invasions of *Ae. albopictus* (Nasci et al. 1989, Lounibos, 2002, Kaplan et al. 2010). The same mechanism was suggested to explain the spread of invasive *Ae. albopictus* in Athens, Greece and the reduction in distribution of the native *Ae. cretinus* (Giatropoulos et al. 2015). Displacement of *Ae. albopictus* by *Ae. aegypti* also occurred in some tropical cities in Asia including Bangkok, Kuala Lumpur, Manilla and southern Taiwan and more recently in the Colombian port of Leticia (reviewed in Lounibos and Juliano, 2018). However, it is suggested that resistance to satyrization may evolve more rapidly in female *Ae. aegypti* populations sympatric to *Ae. albopictus* than in females from allopatric populations (Bargielowski et al. 2013; Lounibos et al. 2016). Bagny et al. (2009) reported a progressive decrease in *Ae. aegypti* distribution on La Réunion island since the 1900s where it was mainly found in rock holes with *Ae. albopictus* in the ravines located on the driest west coast of the island and was absent from artificial containers. The same study suggested that the dwindling *Ae. aegypti* densities observed during the 1950s was due to ecological factors including a competitive interaction between *Ae. aegypti* and *Ae. albopictus* combined with vector control campaigns during these years. However, to the best of our knowledge, the competitive interaction between La Réunion island strains has not yet been investigated. With both *Ae. aegypti* and *Ae. albopictus* being competent vectors of dengue and several other arboviruses including chikungunya and Zika viruses (Lounibos, 2002; Paupy et al. 2009; Chouin-Carneiro et al. 2016), it is important to investigate whether or not the release of irradiated *Ae. albopictus* males would affect *Ae. aegypti* populations (Conseil scientifique de l’Agence française pour la biodiversité 2018), in the framework of the SIT project on La Réunion island. Our satyrization experiments provide insights on the potential impact on *Ae. aegypti* of a control campaign against *Ae. albopictus* on La Réunion island. Different mating crosses were used to test sympatric, conspecific-interspecific and allopatric effects of male *Ae. albopictus* on female *Ae. aegypti* including in a situation of skewed male ratio.

## METHODS

### Mosquito strains, rearing and irradiation conditions

The *Ae. aegypti* strain used in this study originated from field collections on La Réunion island. Colonized in the laboratory by the Institute of Research for Development (IRD) for 5 generations before being transferred to the Insect Pest Control Laboratory (IPCL) of the joint FAO/IAEA division of Nuclear Sciences and Applications, the *Ae. albopictus* La Réunion island strain was then maintained at the IPCL for several generations before these experiments were performed. In order to perform different mating crosses, *Ae. albopictus* strains from China (Guangzhou wild type strain, provided by Wolbaki) and Italy (Rimini strain, provided by Centro Agricoltura Ambiente (CAA)) were maintained in parallel.

All the strains were reared in 30 × 40 × 10 cm trays at a density of 1 first instar larvae (L1) per mL under controlled temperature, humidity and lighting conditions (T= 26± 2 °C, 70± 10 RH%, 12:12h light: dark, including 1h dawn and 1h dusk). Larvae were fed with IAEA larval diet following the protocol described in the Guidelines for routine rearing (IAEA 2017). Pupae were collected and separated using a glass plate sorter (Focks 1980).

Male pupae were irradiated between 36 and 44 hours of age with 40Gy using a Gamma Cell 220 (Nordion Ltd, Kanata, Ontario, Canada) emitting a dose rate of 90Gy/min.

### Experiment 1. Sympatric cross-mating between *Ae. aegypti* and *Ae. albopictus* strains from La Réunion island

Male *Ae. aegypti* and *Ae. albopictus* were irradiated and crossed with female *Ae. albopictus* and *Ae. aegypti*, respectively. Non-irradiated males of each species were also crossed with female mosquitoes of the other species. Non-irradiated male *Ae. aegypti* and *Ae. albopictus* were also crossed with virgin female *Ae. aegypti* and *Ae. albopictus*, respectively, and used as controls.

Three replicates were performed for each cross with 50 males and 50 females transferred to 15 × 15 × 15 cm cages (MegaView Science Co. Ltd., Taiwan) when they were 3 days old for a period of 7 days to ensure enough time was allowed for mating. Females were offered a defibrinated porcine bloodmeal using sausage casings (Grade Specification: 3)26 NC, EDICAS co ltd) for 1 hour (2× 30min with 10min reheating of the blood sausage in between feedings) on two consecutive days when they were 5-6 days old. Each of the females was transferred to an individual drosophila tube containing a cone of seed germination paper (Grade 6, Size: 580 × 580mm, Weight: 145 g/m^2^, Sartorius Stedium Biotech) and 10mL of water. Females were given 3 days to lay eggs and then dissected to determine their insemination status under a stereomicroscope (40 × magnification). Before dissection, females were kept in labelled 50mL Falcon tubes (WWR, Germany) in a refrigerator at 4°C, and samples held in a cold box containing ice to avoid desiccation while other samples were being dissected.

Virgin *Ae. aegypti* and *Ae. albopictus* females of the same batch were offered blood meals and were also placed in individual egging tubes to assess their egging capacity.

All eggs were collected, dried for 7 days in the laboratory and allowed to hatch for 20 h with a hatching solution made of 0.25 g of CM 0001 Nutrient Broth (Oxoid, Hampshire, UK) and 0.05 g of yeast diluted in 0.7 L of deionized water (Zheng et al. 2015).

The number of female mosquitoes still alive after 7 days together with males was recorded for each replicate and survival rate was compared with survival in the conspecific *Ae. aegypti* mating control.

### Experiment 2. Effect of male *Ae. albopictus* density (ratio) on female *Ae. aegypti* mating success

To assess whether an increase in male to female ratio would favour satyrization, female *Ae. aegypti* mosquitoes from La Réunion island were allowed to mate with male *Ae. albopictus* in a male: female ratio of 3:1 corresponding to 75 male *Ae. albopictus* and 25 female *Ae. aegypti*. Three day-old males and females were held in 15 × 15 × 15 cm BugDorm cages (MegaView Science Co. Ltd., Taiwan) for 7 days. The crosses were performed using irradiated males (4 replicates) and non-irradiated males (7 replicates). Females were then dissected to check their insemination status as described above. This experiment was carried out in parallel with Experiment 1 and results of mating crosses could be compared to the insemination rates obtained with a 1:1 male: female ratio (50 males × 50 females).

### Experiment 3. Pre-exposure effect on mating success (interspecific-conspecific treatment)

To assess whether female *Ae. aegypti* that were pre-exposed to male *Ae. albopictus* could re-mate with their conspecific male *Ae. aegypti*, 4 to 5 day-old female *Ae. aegypti* (La Réunion island strain) were pre-exposed to irradiated and non-irradiated male *Ae. albopictus* (La Réunion island strain). Females were removed after 7 days and were offered to irradiated and non-irradiated male *Ae. aegypti* (La Réunion island strain) for another 7 days. Females were then blood fed for 2 consecutive days and eggs collected before females were dissected under a microscope to determine insemination status. We hypothesized that if female *Ae. aegypti* were inseminated by non-irradiated *Ae. albopictus* males, they would not be able to lay fertile eggs even if they had a blood meal.

Fifty adults were included in each cross in a 1:1 male: female ratio in Bugdorm cages (15 × 15 × 15cm) (MegaView Science Co. Ltd., Taiwan).

An experimental design was used which ensured that female *Ae. aegypti* that were pre-exposed to male *Ae. albopictus*, had not been inseminated (Table 1), and female insemination and egg hatch rates were observed.

**Table 1.** Experimental design of the interspecific-conspecific treatments.

| cross | status | number<br>of<br>replicates | male | female | irradiated<br>males | fecundity<br>and fertility<br>checks | dissection for<br>insemination<br>status check |
| --- | --- | --- | --- | --- | --- | --- | --- |
| C1 | control | 5 | AEG | AEG | YES | NO | YES |
| C2 | control | 4 | AEG | AEG | NO | NO | YES |
| C3 | control | 4 | ALBO | ALBO | YES | NO | YES |
| C4 | control | 4 | ALBO | ALBO | NO | NO | YES |
| C5 | pre-exposed | 4 | ALBO | AEG | YES | NO | YES |
| C6 | pre-exposed | 4 | ALBO | AEG | NO | NO | YES |
| C7 | control | 5 | AEG | AEG | NO | YES | YES |
| C8 | pre-exposed | 5 | ALBO | AEG | NO | YES | YES |

AEG and ALBO stand for *Ae. aegypti* and *Ae. albopictus* respectively; C for cross and numbers 1-8 for cross identity

### Experiment 4. Effect of geographic origin on mating success (allopatric crosses)

Since we observed resistance to satyrization against *Ae. albopictus* in the sympatric cross experiment, we explored the allopatric effect to better understand the mating behavior of the La Réunion island strain of *Ae. aegypti*. Several crosses were performed using irradiated and non-irradiated males of the *Ae. albopictus* strains from La Réunion island, China (Guangzhou wild type strain) and Italy (Rimini strain).

Female *Ae. aegypti* mosquitoes from La Réunion island were crossed for 7 days with non-irradiated or irradiated male *Ae. albopictus* mosquitoes from either La Réunion island, China or Italy in 14 combinations. In each interspecific treatment (*Ae. aegypti* female × *Ae. albopictus* male), 30 males and 30 females, 3 days old, from each strain were housed in 15 × 15 × 15 cm BugDorm cages (MegaView Science Co. Ltd., Taiwan). In addition, conspecific control crosses (*Ae. albopictus* female × *Ae. albopictus* male, and *Ae. aegypti* female × *Ae. aegypti* male, La Réunion island strain) were carried out using the same number of males and females for each strain. The number of females that were successfully inseminated was recorded (as described for Experiment 1) and results were compared to the interspecific crosses and to the conspecific control crosses. Egg hatch was performed as described for Experiment 1 for batches that were collected from females mated with non-irradiated males.

### Statistics

Statistical analyses were performed and graphs drawn using RStudio Team (2016). The proportion of inseminated females was calculated as the number with at least one spermathecae filled with sperm divided by the number of dissected females. The proportion egg hatch was calculated from the number of hatched eggs divided by the total number of eggs laid per individual female (Experiment 1) or per group of females in a cage (Experiments 3 and 4). The proportions were transformed following the Freeman–Tukey method (arcsine transformed data). An analysis of variance (ANOVA) was then performed followed by a Tukey multiple comparison to compare means of each crossed pair. A paired t-test was performed to compare egg hatch between pre-exposed and non-exposed females (Pre-exposure effect on mating success experiment).

The survival rate (Experiment 1) was analyzed using a generalized binomial linear mixed-effects model fit by maximum likelihood (Laplace approximation) with logit link with the survival rate, defined as dependent variable, and type of mating pairs (cross) as fixed effects and replicates as a random effect. The best model was selected based on the lowest corrected Akaike information criterion (AICc), and the significance of fixed effects was tested using the likelihood ratio test (Burnham and Anderson, 2003; Hurvich and Tsai, 1995).

The statistical significance for all experiments was determined at the α = 0.05 level.

## RESULTS

### Experiment 1. Sympatric cross-mating between *Ae. aegypti* and *Ae. albopictus* strains from La Réunion island

There was no insemination between sympatric species from La Réunion island strains either when non-irradiated or irradiated male *Ae. aegypti* or *Ae. albopictus* were caged with female *Ae. albopictus* (n=82, mean (± SE) =0±0% and n=72, mean (± SE) =0±0%) or *Ae. aegypti* (n=28, mean (± SE) =0±0% and n=41, mean (± SE) =0±0%), respectively, compared to the conspecific mating controls (*Ae. albopictus*: n= 58, mean (± SE) =99.2±0.01%; *Ae. aegypti*: n=71, mean (± SE) =100±0%) (Figure 1, F _(5,12)_ = 735.6, *p* < 0.0001). A Tukey multiple comparisons of means did not show any difference in insemination rate regardless of the cross type between irradiated or non-irradiated males of *Ae. albopictus* or *Ae. aegypti* and non-irradiated females of *Ae. aegypti* and *Ae. albopictus*, respectively (*p* > 0.05).

**Figure 1.**
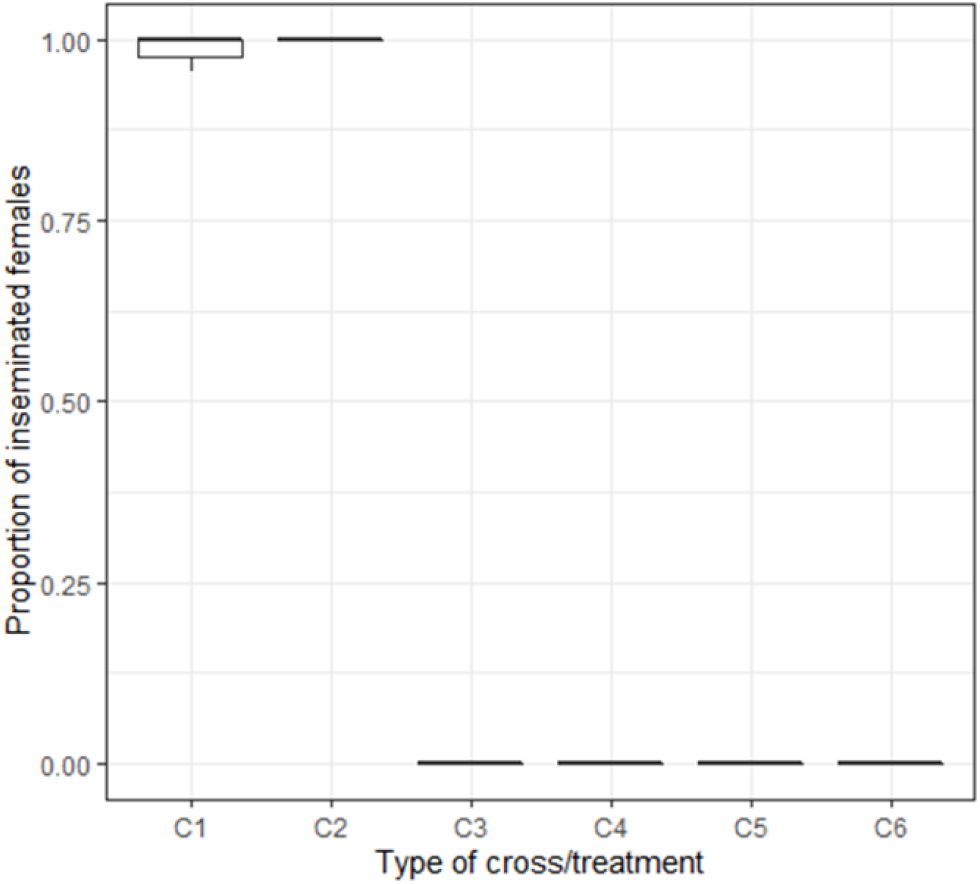
Sympatric cross-mating between *Ae. aegypti* and *Ae. albopictus* strains from La Réunion island. C denotes the cross and numbers (1-6) are related to the cross identity. The number of dissected females (n) per cross were C1 =control *Ae. albopictus* (non-irradiated), n=58), C2= control *Ae. aegypti (*non-irradiated), n=71, C3= male *Ae. albopictus* (non-irradiated) × female *Ae. aegypti*, n=28, C4= male *Ae. albopictus* (irradiated) × female *Ae. aegypti*, n=41, C5= male *Ae. aegypti* (non-irradiated) × female *Ae. albopictus*, n=82, C4= male *Ae. aegypti* (irradiated) × female *Ae. albopictus,* n=72.

Regardless of the male status (irradiated or non-irradiated) the females exposed to interspecific mating, and virgin females, laid a lower number of eggs per female than the females of the conspecific mating control crosses (Interspecific: (C3-6): 146 eggs (n=223); virgin females (Faeg-Falbo): 92 eggs (n=77), controls (C1-2): 2 047eggs (n=129)). None of eggs from interspecific mating crosses and virgin females hatched compared to the controls (Figure 2, F _(7, 46)_ =84.73, *p* < 0.0001). Interspecific mating was more detrimental to female *Ae. aegypti* survival than to *Ae. albopictus* (Table 2, *p*<0.0001).

**Table 2.** Comparison of survival rate between female *Ae. aegypti* in the cross-mating between sympatric *Ae. aegypti* and *Ae. albopictus* strains from La Réunion island.

| | Estimate | Std. Error | z value | $p(>\vert z\vert )$ |
| --- | --- | --- | --- | --- |
| (Intercept) | 5.93E-16 | 1.63E-01 | 0 | 1 |
| crossctrlalbo | -2.96E-01 | 2.32E-01 | -1.272 | 0.2032 |
| crossMaegFalbo | 5.04E-01 | 2.02E-01 | 2.492 | 0.0127 * |
| crossMalboFaeg | -1.05E+00 | 2.10E-01 | -4.987 | 6.13e-07 *** |

Std. standard
Ctrlalbo stands for control *Ae. albopictus* cross; MaegFalbo=male *Ae. aegypti* × female *Ae. albopictus*;
MalboFaeg=male *Ae. albopictus* × female *Ae. aegypti*

**Figure 2.**
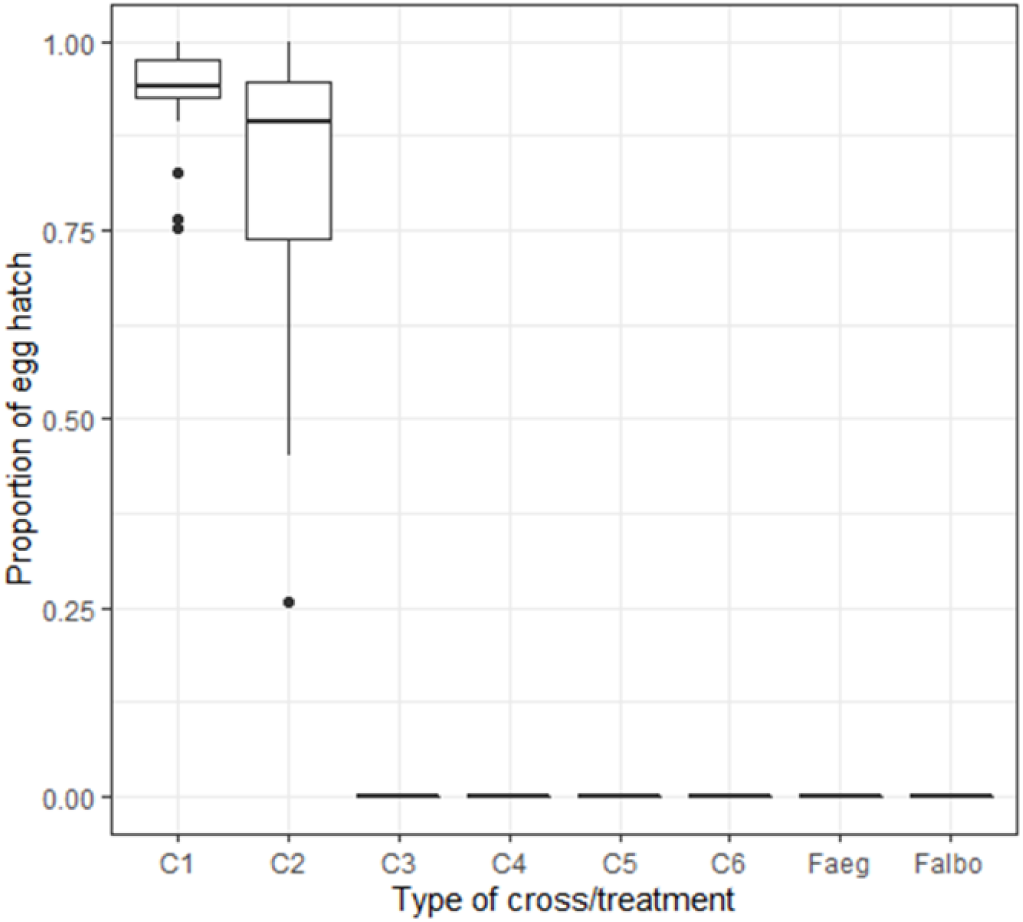
Egg hatch in sympatric crosses between *Ae. aegypti* and *Ae. albopictus* strains from La Réunion island. C denotes the cross and numbers (1-6) are related to the cross identity. C1 =control *Ae. albopictus* (non-irradiated), C2= control *Ae. aegypti (*non-irradiated), C3= male *Ae. albopictus* (non-irradiated) × female *Ae. aegypti*, C4= male *Ae. albopictus* (irradiated) × female *Ae. aegypti*, C5= male *Ae. aegypti* (non-irradiated) × female *Ae. albopictus*, C4= male *Ae. aegypti* (irradiated) × female *Ae. albopictus.*

### Experiment 2. Effect of male *Ae. albopictus* density (ratio) on female *Ae. aegypti* mating success

Increasing the ratio of male: female to 3:1 in favour of male *Ae. albopictus* did not significantly increase the satyrization effect on female *Ae. aegypti* (La Réunion island) judged by insemination rates for both non-irradiated (t-test, ratio 1:1: mean (± SE) =0±0%, n=28, ratio 3:1: mean (± SE) =2±0.1%, n=169, *p*=0.26) and irradiated (t-test, ratio 1:1: mean (± SE) =0±0%, n=41, ratio 3:1: mean (± SE) =0±0 %, n=101, *p*=1) male *Ae. albopictus.* All these crosses had negligible insemination rates compared to conspecific mating (controls) (F _(7, 21)_ = 272.8, *p*<0.0001).

### Experiment 3. Pre-exposure effect on mating success

When female *Ae. aegypti* were pre-exposed to non-irradiated or irradiated male *Ae. albopictus*, there was no insemination (Figure 3. C5: mean (± SE) =0±0%, n=90, and C6: mean (± SE) =0±0%, n=121) whereas groups of females that were pre-exposed to male *Ae. albopictus* were inseminated by their male *Ae. aegypti* conspecifics (Figure 3. C7: mean (± SE) =100±0%, n=106, and C8: mean (± SE) =95.99±0%, n=146). No difference in female insemination rates was observed between pre-exposed and non-exposed females (*p*=0.27) or between pre-exposed females and controls (Figure 3. C1-3 vs C7-8) (*p* > 0.05).

**Figure 3.**
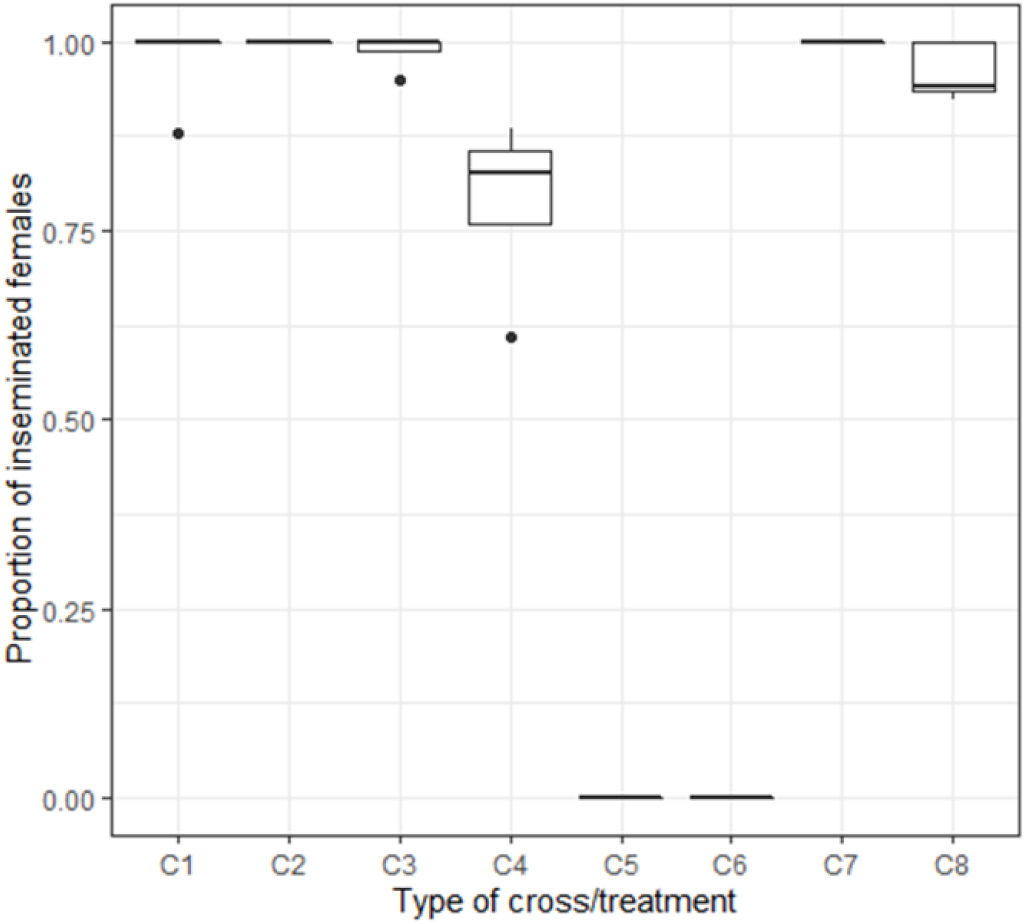
Pre-exposure effect on mating success. C denotes the cross and numbers (1-8) are related to the cross identity. C1 = control *Ae. aegypti* (non-irradiated), C2 = control *Ae. aegypti (*irradiated), C3 = control *Ae. albopictus* (non-irradiated), C4 = control *Ae. albopictus* (irradiated), C5 = male *Ae. albopictus* (non-irradiated) × female *Ae. aegypti*, C6 = male *Ae. albopictus* (irradiated) × female *Ae. aegypti*, C7 = male *Ae. aegypti* (non-irradiated) × female *Ae. aegypti (*non-exposed to male *Ae. albopictus)*, C8 = male *Ae. aegypti* (non-irradiated) × female *Ae. aegypti* (pre-exposed to male *Ae. albopictus*).

Female *Ae. aegypti* that were pre-exposed to male *Ae. albopictus* successfully laid fertile eggs when they then mated with their male *Ae. aegypti* conspecifics, and mean egg hatch was lower but not significantly different between the non-exposed (mean (± SE) =95.99±0.008%) and pre-exposed females (mean (± SE) =91.61±0.02%) (t-test: t=2.0576, df=4, *p*=0.1).

### Experiment 4. Effect of geographic origin on mating success (allopatric crosses)

A greater female insemination rate was observed when conspecific mating was compared to the interspecific mating groups, regardless of the male origin or irradiation status (Table 3, F _(13, 23)_ = 229.1, *p*<0.0001). Neither geographic origin nor irradiation status of male *Ae. albopictus* had an impact on female *Ae. aegypti* insemination rate, confirming the conclusion that allopatric and sympatric effects were similar when the La Réunion island strain of *Ae. aegypti* was used in these experiments (Table 3, *p*>0.05).

**Table 3.** Mean insemination and egg hatch rates (±SE) in crosses between female *Ae. aegypti* (La Réunion island strain) and *Ae. albopictus* males from China, Italy (allopatric) and La Réunion island (sympatric).

| Mating type | Male |  | Female | Insemination rates (%) | hatch rates (%) |
| --- | --- | --- | --- | --- | --- |
|  | irradiated | non-irradiated |  |  |  |
| interspecific | China | | Réunion | 0.92±0.92<sup>a</sup> (95) | NA |
| | | China | Réunion | 0±0<sup>a</sup> (82) | 0±0<sup>a</sup> |
| conspecific | China | | China | 100±0<sup>b</sup> (24) | NA |
| | | China | China | 100±0<sup>b</sup> (48) | 100±0<sup>b</sup> |
| interspecific | Réunion | | Réunion | 1.92±1.11<sup>a</sup> (94) | NA |
| | | Réunion | Réunion | 1.33±1.33<sup>a</sup> (74) | 0±0<sup>a</sup> |
| conspecific | Réunion | | Réunion | 100±0<sup>b</sup> (57) | NA |
| | | Réunion | Réunion | 100±0<sup>b</sup> (54) | 71.8±2.7<sup>c</sup> |
| interspecific | Italy | | Réunion | 0±0<sup>a</sup> (99) | NA |
| | | Italy | Réunion | 5.5±3.68<sup>a</sup> (77) | 0±0<sup>a</sup> |
| conspecific | Italy | | Italy | 100±0<sup>b</sup> (50) | NA |
| | | Italy | Italy | 100±0<sup>b</sup> (57) | 83.5±1.55<sup>c</sup> |
| conspecific | Réunion* | | Réunion | 100±0<sup>b</sup> (72) | NA |
| | | Réunion* | Réunion | 100±0<sup>b</sup> (54) | 89.8±2.11<sup>c</sup> |

‘Interspecific’ stands for crosses between female *Ae. aegypti*, La Réunion island strain and *Ae. albopictus* (strain from China, Italy, La Réunion island) and ‘conspecific’ for control mating between male and female of the same species. Réunion\*= male *Ae. aegypti* from La Réunion island. Numbers into parentheses represent the number of dissected females for insemination assessment. Different letters show significant difference between groups.

We observed a significantly lower egg hatch in interspecific than conspecific crosses irrespective of the male origin (Table 3, F _(13, 23)_ = 182.9, *p*<0.0001).

## DISCUSSION

The objective of this study was to assess the level of reproductive interference between male *Ae. albopictus* and female *Ae. aegypti* and to discuss the strategy of management of the two vector species in a sympatric area. In laboratory conditions almost no satyrization effect was observed between sympatric populations of *Ae. aegypti* and *Ae. albopictus* colonised from La Réunion island, even when the male *Ae. albopictus* to female *Ae. aegypti* ratio was increased to 3:1. Female *Ae. aegypti* previously mated to male *Ae. albopictus* were not prevented from being inseminated by conspecific male *Ae. aegypti*. Satyrization did not occur between allopatric *Ae. albopictus* and *Ae. aegypti* strains from La Réunion island either. An *Ae. aegypti* strain from La Réunion island has therefore developed full resistance to satyrization (anti-satyrization behavior).

Many reasons could explain the observed low level of satyrization. Bargielowki et al. (2013; 2015) have previously described sympatric field populations that have co-existed long enough to evolve resistance to cross-mating and showed that *Ae. aegypti* females allopatric to *Ae. albopictus* are more susceptible to interspecific insemination by *Ae. albopictus* (Lounibos et al. 2016). Cage experiments and field observations indicate that *Ae. albopictus* males are capable of satyrizing females of other species of the Stegomyia subgenus, potentially leading to competitive displacement, and even extinction, especially of endemic species on islands (Bargielowki et al. 2015). In contrast, in our study even an increase in ratios in favour of male *Ae. albopictus* did not significantly increase insemination of female *Ae. aegypti*. One study has pointed to the implication of population density on mating behavior (Marcela et al. 2015), which found that male density significantly increased swarming behavior, number of mating pairs, and egg production of hetero-specific females, but not female insemination. They also found that *Ae. aegypti* males mate more readily with hetero-specific females than do *Ae. albopictus* males and so if *Ae. aegypti* were released into the field they may mate with both *Ae. aegypti* and *Ae. albopictus* females, and reduce populations of both species by preventing offspring. There is no evidence that this would be the case on La Réunion island since we observed low reproductive success when crosses were performed in both directions. In addition, we observed that interspecific mating was detrimental to female *Ae. aegypti* survival. It has been previously documented that interspecific copulation and hybridization can reduce female reproductive success, but interspecific sexual harassment, which does not always result in interspecific copulation, can also adversely affect individual reproductive success and longevity by suppressing both sugar and blood feeding (Soghigian et al. 2014). In white butterflies (*Leptidea* spp.), for example, the prolonged mating ritual of hetero-specific males decreases the number of opportunities that females have to mate with conspecific males even when interspecific copulation does not take place (Friberg et al. 2013).

Similarly, in bean weevils (*Callosobruchus* spp.), males persistently chase hetero-specific females, causing reductions in the oviposition rate and shortened longevity of these females (Kishi et al. 2009). *Ae. aegypti* females pre-exposed to male *Ae. albopictus* were able to be inseminated by their conspecific male *Ae. aegypti* demonstrating that the La Réunion island *Ae. aegypti* strain has developed a resistance to satyrization. Carrasquilla and Lounibos (2015) have showed that *Ae. aegypti* females, previously exposed to *Ae. albopictus* males, were rendered refractory to subsequent conspecific mating even when their spermathecae contained no hetero-specific sperm. Additional experiments demonstrated transfer of labelled semen from *Ae. albopictus* males to *Ae. aegypti* females and low production of viable eggs of females housed with conspecific males, following exposure to *Ae. albopictus* males, and confirmed higher incidence of satyrization than expected, based on hetero-specific insemination rates. We did not observe this result after pre-exposing *Ae. aegypti* females to *Ae. albopictus* for 7 days before replacing male *Ae. albopictus* by male *Ae. aegypti* mosquitoes. It has been shown that interspecific pairs needed more time together before mating occurred. Bargielowki et al. (2015) found that when female *Ae. aegypti* were exposed for up to 3 weeks, interspecific insemination increased significantly from 1 % after 1 day, to 10% after 1 week and to more than 50% after 3 weeks. However, assuming that most released sterile males will survive around 1 week in the field, it is unlikely that most released males would be able to find and mate with females after 3 weeks in the wild (Bellini et al. 2010; Iyaloo et al. 2019). These results indicate that in areas where *Ae. aegypti* and *Ae. albopictus* co-occur, releasing sterile male *Ae. albopictus* may not suppress *Ae. aegypti* populations. It would be more beneficial to suppress the species with the smallest population, before further planning to control the second species, assuming the epidemiological impact of each species was equal.

In our study, female *Ae. aegypti* pre-exposed to male *Ae. albopictus* produced eggs which had similar egg hatch when re-mated with their conspecifics, meaning that females had not been inseminated by the *Ae. albopictus* males. Those females which later mated with their conspecifics and laid eggs were apparently fully fertilized by conspecific sperm. However, it has been shown that the satyrization effect could be underestimated when evaluation of mating status of females is based on whether the spermathecae were filled with sperm or not (Carrasquilla and Lounibos, 2015). It therefore cannot be ruled out that some females might have been inseminated when pre-exposed to *Ae. albopictus* based on the variation observed in egg hatch. Bargielowski et al. (2015) demonstrated that multiple inseminations can occur in older female *Ae. aegypti* when the effects of accessory gland proteins have worn off, and in females mated to sperm-depleted males. In any case, hetero-specific sperm is known to be stored in separate spermathecae (Bargielowski et al. 2015) and so was presumably not significantly used for egg fertilization.

Allopatric *Ae. albopictus* males did not perform better than sympatric males and anti-satyrization effects seem to protect against allopatric populations. This shows that resistance to one strain confers resistance to others. Honórios et al. (2018) demonstrated that only some populations of *Ae. albopictus* are capable of satyrization. Female *Ae. aegypti* from populations allopatric to *Ae. albopictus* in the field were more susceptible to interspecific mating than females from sympatric populations, and selection experiments in cages confirmed the rapid development of resistance to satyrization in the laboratory, as well as changes in behavior toward conspecifics associated with increased satyrization resistance (Bargielowski and Lounibos 2014). The fact that the *Ae. aegypti* populations persist in La Réunion island ravines as opposed to urban environments could be due to some genetic differentiation from domestic subspecies. Lounibos and Juliano (2018) have recently pointed out that the feral subspecies *Ae. aegypti formosus* is expected to behave differently than the domestic subspecies but populations of this species from Madagascar, La Réunion island and Mayotte have not been tested yet for genetic distinctiveness from *Ae. aegypti* (*aegypti)* to the best of our knowledge. In any case, a signature of selection in the *Ae. aegypti* genome to a specific type of interspecific interaction (mating) was found by Reiskind et al. (2017) allowing the identification of its genetic basis.

When considering a regional approach for *Aedes* control using the SIT, compatibility of strains as well as species may be important as it would allow strains to be imported for release from nearby countries where they can be more easily reared and/or irradiated. Damiens et al. (2016) demonstrated that male *Ae. albopictus* from Mauritius and Seychelles islands, about 50-200 km away from La Réunion island, were compatible and could successfully inseminate female *Ae. albopictus* regardless of their origin. A regional SIT mass-rearing programme could therefore be envisaged, with a good transportation method, but the release of sterile *Ae. albopictus* males may not have the added benefit of satyrizing the local *Ae. aegypti* population if an overflooding of irradiated male *Ae. albopictus* leads to similar results.

The development of resistance to satyrization in the *Ae. aegypti* strain shows that strong competition between the sympatric *Ae. aegypti* and *Ae. albopictus* probably occurs on La Réunion island. An SIT project against *Ae. albopictus* would not have an effect on *Ae. aegypti* populations, and other mechanisms such as larval competition probably explain the current geographical retraction of *Ae. aegypti* to the ravines. Bagny et al. (2013) observed that this narrow distribution of *Ae. aegypti* was due to its poorer ability to cope with unfavourable temperatures and to its lower competition between larvae for resources compared to *Ae. albopictus* (Juliano 2009). The two species may co-exist as long as the dominant *Ae. albopictus* is present and the resistance could be maintained by satyrization pressure (Bargielowski et al. 2019). Global climate change may favour an increase in the population size of *Ae. aegypti* (Juliano et al. 2004) which is a greater vector of arboviral diseases including dengue, chikungunya, yellow fever and Zika. Therefore, suppressing or eliminating *Ae. albopictus* will likely promote expansion of *Ae. aegypti* (HCB Scientific Committee 2017). Whilst it may be important to target the most epidemiologically important vector first (Alphey et al. 2010), considering its limited distribution, the eradication of *Ae. aegypti* population may be seen as a first priority.

